# Phthiocerol dimycocerosates from *Mycobacterium tuberculosis* increase the membrane activity of bacterial effectors and host receptors

**DOI:** 10.1101/2020.05.13.092585

**Authors:** Jacques Augenstreich, Evert Haanappel, Fadel Sayes, Roxane Simeone, Valérie Guillet, Serge Mazeres, Christian Chalut, Lionel Mourey, Roland Brosch, Christophe Guilhot, Catherine Astarie-Dequeker

**Affiliations:** Institut de Pharmacologie et de Biologie Structurale (IPBS), Université de Toulouse, CNRS, UPS, F-31077 Toulouse, France; Institut Pasteur, Unit for Integrated Mycobacterial Pathogenomics, CNRS UMR3525, F-75724 Paris, France

**Keywords:** Macrophages, tuberculosis, *Mycobacterium*, phthiocerol dimycocerosates, EsxA, membranolytic activity, receptors, membranes, complement receptor 3

## Abstract

*Mycobacterium tuberculosis* (*Mtb*) synthesizes a variety of atypical lipids that are exposed at the cell surface and help the bacterium infect macrophages and escape elimination by the cell’s immune responses. In the present study, we investigate the mechanism of action of one family of hydrophobic lipids, the phthiocerol dimycocerosates (DIM/PDIM), major lipid virulence factors. DIM are transferred from the envelope of *Mtb* to host membranes during infection. Using the polarity-sensitive fluorophore C-Laurdan, we visualized that DIM increase the membrane polarity of a supported lipid bilayer put in contact with mycobacteria, even beyond the site of contact. We observed that DIM activate the complement receptor 3, a predominant receptor for phagocytosis of *Mtb* by macrophages. DIM also increased the activity of membrane-permeabilizing effectors of *Mtb*, among which the virulence factor EsxA. This is consistent with previous observations that DIM help *Mtb* disrupt host cell membranes. Taken together, our data show that transferred DIM spread within the target membrane, remodel lipid organization and increase the activity of host cell receptors and bacterial effectors, diverting in a nonspecific manner host cell functions. We therefore bring new insight into the molecular mechanisms by which DIM increase *Mtb’s* capability to escape the cell’s immune responses.

## Introduction

The envelope lipids, phthiocerol dim*y*cocerosates (DIM/PDIM) are among the major virulence factors of *Mycobacterium tuberculosis* (*Mtb*). Apart from their structural role as components of the mycobacterial envelope, DIM are harnessed by the bacteria to manipulate host immune functions, especially during the early steps of infection, when *Mtb* encounters macrophages (Rousseau et al., 2004;Astarie-Dequeker et al., 2009;Cambier et al., 2014;Passemar et al., 2014;Barczak et al., 2017;Quigley et al., 2017). However, there is still a great deal of uncertainty about the molecular mechanisms of DIM-mediated effects in host cells.

Our group has reported that DIM are transferred from the mycobacterial envelope to the membranes of host macrophages, and that this transfer increases the phagocytosis of bacteria (Augenstreich et al., 2019). Further investigations indicated that once inserted in the host membrane, DIM adopt a conical shape which disturbs the organization of membrane lipids (Augenstreich et al., 2019). We hypothesize that this has functional consequences for the activity of membrane-associated contributors to phagocytosis. Phagocytosis occurs through the recognition of Pathogen-Associated Molecular Patterns (PAMPs) present at the bacterial surface by receptors expressed in the host cell plasma membrane. In the case of *Mtb*, phagocytosis involves multiple receptors, including the complement receptor 3 (CR3, also named CD11b/CD18 or Mac-1) (Schlesinger, 1993;Stokes et al., 1993;Cywes et al., 1996). Interestingly, blocking CR3 with antibodies significantly reduces the entry of a DIM-proficient *Mtb* strain into macrophages without affecting the uptake of a DIM-deficient mutant (Astarie-Dequeker et al., 2009), suggesting that the disorganizing effects of DIM on host cell membranes could target the activity of CR3.

DIM also cause membrane damage, such as phagosomal rupture and cell death (Augenstreich et al., 2017;Quigley et al., 2017). Previous work has shown that DIM act in concert with the type VII secretion system ESX-1 (T7SS/ESX-1), in a manner that involves the 6-kDa early secreted antigenic target (ESAT-6, also known as EsxA) (Augenstreich et al., 2017;Barczak et al., 2017). EsxA has been predicted to insert into membranes and to lyse artificial and biological membranes (Hsu et al., 2003;Gao et al., 2004;de Jonge et al., 2007;Smith et al., 2008;Ma et al., 2015;Peng et al., 2016). Interestingly, the membranolytic activity of EsxA was found to depend on the lipid composition and on membrane fluidity (Augenstreich et al., 2017;Ray et al., 2019). However, the mechanism by which DIM and EsxA collaborate remains unclear. Barczak and colleagues proposed that the secretion of EsxA requires DIM (Barczak et al., 2017). However, previous findings indicated that the DIM-deficient mycobacterial strains, which are affected in their membranolytic activity, secrete similar amounts of EsxA as the parental strains (Augenstreich et al., 2017;Quigley et al., 2017). Recently, our group reported that the incorporation of DIM in the membrane of model liposomes enhanced EsxA activity (Augenstreich et al., 2017). Nevertheless, the origin of the membrane-disrupting activity of DIM remains obscure. Indeed, their potentiating effect on EsxA’s activity is independent of their ability to rigidify membranes and is limited to membranes containing lipids extracted from macrophages (Augenstreich et al., 2017). Moreover, the membranolytic activity of EsxA itself was called into question by Conrad and colleagues who showed that the hemolytic activity of recombinant EsxA obtained from the same source was due to residual detergent used in the purification protocol (Conrad et al., 2017). Thus, the capacity of *Mtb* to permeabilize host membranes is more complex than initially thought.

It therefore remains of importance to determine how DIM contribute to the phagocytosis of *Mtb* and to the membranolytic activity of bacteria at the molecular level.

## Material and Methods

### Antibodies and reagents

The mouse monoclonal antibody directed against the extracellular domain of human CR3, clone 2LPM19c) (IgG1, dilution 1:20) and normal mouse IgG1 were purchased from Santa Cruz Biotechnology. The CBRM1/5 mouse antibody raised against an activation-specific epitope on the CD11b subunit of human CR3 (IgG1, dilution 1:50) was from eBiosciences. POPC (1-palmitoyl-2-oleoyl-sn-glycero-3-phosphocholine) was from Avanti Polar Lipids. THP-1 lipids were extracted from THP-1 cells as described previously (Augenstreich et al., 2017). C-Laurdan was synthesized as described in (Mazeres et al., 2014). DIM (m_w_ = 1432 g/mol) were purified from *M. canetti* (Augenstreich et al., 2019). Stock solutions of 40 mg/mL were prepared by dissolving the dried lipids in CH_3_OH/CHCl_3_ (2:1, vol/vol) (Augenstreich et al., 2019). For cell treatment, DIM were dispersed in culture medium at a final concentration of 0.1 mg/mL corresponding to 70 μM. The other reagents were purchased from Sigma-Aldrich.

### Bacterial strains and growth conditions

The unmarked *mas* mutant (H37RvΔ*mas*), the *ppsE* mutant (H37Rv Δ*ppsE*), the *esxA* mutant (H37RvΔ*esxA*) of *Mtb*, the *mas* mutant (BCGΔ*mas*), the spontaneous *fad26* mutant (BCGΔ*fad26*) and the *pks15/1 mutant* (BCGΔ*pks15/1)* of *M. bovis* BCG *and* the recombinant BCG strains complemented with the ESX-1 system in DIM-proficient or DIM-deficient strains (BCG::ESX-1 and BCGΔ*mas*::ESX-1) were described in previous work (Constant et al., 2002;Astarie-Dequeker et al., 2009;Simeone et al., 2010;Augenstreich et al., 2017). Bacteria were rendered fluorescent by transferring the plasmid pMV361H *gfp* (Astarie-Dequeker et al., 2009) or the plasmid pMVmCherry derived from a plasmid pMV361eH harbouring the mCherry encoding gene (Burbaud et al., 2016). All strains were cultured at 37°C as previously described (Augenstreich et al., 2017)

### Phagocyte culture

The human promonocytic cell line THP-1 (ECACC 88081201; Salisbury, UK) was cultured and differentiated in macrophages with PMA, as mentioned in (Augenstreich et al., 2019). Human blood purchased from the Etablissement Français du Sang (Toulouse, France) was collected from fully-anonymous non-tuberculous donors. Peripheral blood mononuclear cells (PBMC) and human monocyte-derived macrophages (hMDM) were prepared as previously described (Astarie-Dequeker et al., 2009).

### Phagocytosis assay

Single bacteria suspensions were prepared from exponentially growing strains as previously described (Astarie-Dequeker et al., 2009;Tabouret et al., 2010). The concentration of bacteria was estimated from the optical density (OD) at 600 nm. hMDM cultured in RPMI on glass coverslips were incubated at 37°C for 1 h with GFP-expressing bacteria at a multiplicity-of-infection (MOI) of 10:1 or with zymosan at MOI 30:1. When indicated, cells were pre-treated for 30 min with 10 μg/mL anti-CR3 blocking antibody or non-relevant IgG1 and for another 1 h with 70 μM DIM or equivalent volume of solvent. Phagocytosis was assessed by fluorescence microscopy as previously reported (Astarie-Dequeker et al., 2009;Augenstreich et al., 2019).

### Expression and purification of recombinant EsxA

Recombinant *Mtb* EsxA (rEsxA) was prepared according to the step-by-step protocol provided by BEI Resources Repository (available at csu-cvmbs.colostate.edu/Documents/dobos-rp004.pdf). The C-terminally His-tagged protein was expressed in *Escherichia coli* BL21(DE3)pLysS using the plasmid pMRLB.7 containing *Mtb esxA* (BEI Resources). rEsxA was then purified according to the BEI protocol, adapted by omitting the washing step with ASB14. The protein was checked for purity by SDS/PAGE followed by Coomassie staining, and dialyzed using 3,500-MWCO dialysis tubing in 10 mM ammonium bicarbonate, lyophilized and stored at −20°C. For the calcein leakage experiments, a small amount of rEsxA was weighed and solubilized in PBS at a final concentration of 2 mg/mL.

### Assay for measurement of membrane polarity

We characterized membrane polarity using the membrane dye C-Laurdan, whose fluorescence spectrum shifts toward lower wavelengths when the membrane becomes more apolar (Mazeres et al., 2017). A supported lipid membrane of phospholipid POPC labeled with 1% C-Laurdan was formed on a Ø25 mm microscope coverslip using the method of vesicle fusion (Mascalchi et al., 2012). The membrane was put in contact with a suspension of 2 × 10^6^ bacteria, centrifuged at 1,500g for 5 min at room temperature to sediment the bacteria onto the membrane, and further incubated for 20 min at room temperature. We took two-photon microscopy images on a Zeiss LSM710 inverted confocal microscope using a 40X water immersion objective. The membrane polarity was characterized by the Generalized Polarization (GP), a ratiometric measure given by GP = (*I_440_* – *I_490_*)/ (*I_440_* + *I_490_*) where *I_440_* and *I_490_ are* the fluorescence intensities of C-Laurdan at 440 nm and 490 nm, respectively. The value of GP increases when the membrane becomes more apolar.

### Calcein leakage assays

The membranolytic activity of compounds was evaluated on liposomes using a calcein leakage assay, as previously described (Augenstreich et al., 2017). The liposomes were composed of POPC or of THP-1 lipids supplemented or not with 10% (mol/mol) of DIM and contained a solution of 50 mM calcein.

### Contact dependent-hemolytic activity of *Mycobacterium spp*

Lytic activity of *Mycobacterium spp* was detected by an hemolysis assay. Exponentially-growing mycobacteria in 7H9-ADC-Tween 80 0.05% were pelleted by centrifugation and washed twice in PBS (Smith et al., 2008;Conrad et al., 2017). The concentration of bacteria in suspension was estimated from the OD at 600 nm. Fresh blood was centrifuged and the pellet was resuspended in isotonic NaCl solution and washed twice with PBS. The final red blood cells (RBC) suspension containing 1 × 10^7^ cells was mixed with bacteria at MOI of 50:1, centrifuged and incubated at 37°C for 48 h. The pellet was resuspended, centrifuged and the supernatant was recovered. The absorbance at 415 nm (A_415_) was measured on a CLARIOstar 96-well microplate reader. The supernatants of cells treated with PBS or lysed with 0.1% Triton X-100 were used as reference values for 0% and 100% hemolysis, respectively. The percentage of hemolysis was calculated using the formula: %hemolysis = (A_415,sample_ – A_415,PBS_) / (A_415,TritonX100_ – A_415,PBS_).

### Statistics

All results are expressed as mean ± standard error of the mean (SEM) for the indicated number of experiments (n). Statistical analyses were performed using GraphPad Prism 6.0 (GraphPad Software Inc.) and presented in Figure legends. Differences were considered significant if p<0.05.

## Results

### DIM contribute to the membrane polarity decrease in a lipid bilayer beyond the site of interaction with mycobacteria

We previously reported that DIM are transferred from *M. bovis* BCG to the macrophage (Augenstreich et al., 2019) and decrease its overall membrane polarity (Astarie-Dequeker et al., 2009). We investigated whether this decrease is restricted to the site of contact between the bacilli and the host membrane, or extends beyond it, as the pleiotropic effects of DIM suggest. A supported POPC membrane labelled with the polarity-sensitive fluorophore C-Laurdan was put in contact with mCherry-expressing bacilli, either wild-type *M. bovis* BCG (BCG WT) or a DIM-deficient isogenic *M. bovis* BCG mutant, BCGΔ*mas*, with a deleted *mas*, encoding an enzyme essential for DIM synthesis (Azad et al., 1996). Using mCherry fluorescence and transmission images, we first searched for bacteria immobilized on the membrane (Figure 1a, left). The majority of bacteria remained in suspension and diffused in the buffer (blurred spots in Figure 1a). By two-photon microscopy, we then imaged the C-Laurdan around immobile bacteria. Bacteria absorbed C-Laurdan from the bilayer (Figure 1a, right), implying that the C-Laurdan fluorescence spectrum at the point of contact contained contributions both from the membrane and the bacteria. We discerned no difference in C-Laurdan fluorescence intensity between the two strains (Figure S1a). We enlarged the bacterial contour, determined from the C-Laurdan fluorescence, three times by 5 pixels (520 nm), thus defining three concentric bands around the immobile bacterium (C1, C2 and C3 in Figure 1b image). From the average fluorescence spectrum in these bands, we calculated the corresponding Generalized Polarisation (GP) values to characterize the membrane polarity in each band. We subtracted the intrinsic GP of the membrane, averaged over three zones far from the bacteria, from the GP of the different bands to quantify the GP difference (ΔGP) around bacteria. GP was increased inside the bacterial contour for both strains, reflecting the apolar nature of the mycobacterial envelope. We noticed a smaller increase in GP in zone C1 with respect to the membrane for both strains (Figure S1b). However, further away from the bacteria in band C2 and C3, ΔGP remained positive for BCG WT, whereas GP returned to the membrane reference value for BCGΔ*mas* (Figure S1b). Hence, the mean GP increase was both higher and observed over a larger distance for the DIM-producing BCG WT strain than for the DIM-deficient BCGΔ*mas* mutant (Figure 1b). The membrane was therefore more apolar around BCG WT than around BCGΔ*mas*. Given the resolution of the microscope (around 250 nm), the observed GP increase in C1 might be caused by bacterial fluorescence extending into C1. Importantly, this increase persisted up to zone C3 for BCG WT corresponding to 1-1.5 μm from the edge of the bacteria (Figure 1b). This distance is too big to be caused by an optical resolution effect. In *M. bovis* BCG, the deletion of *mas* induces not only DIM deficiency but also the loss of the structurally similar phenolic glycolipids (PGL). Therefore, we tested BCG mutants deficient for DIM or PGL only. The BCGΔ*fad26* mutant, having a spontaneous mutation in the *fad26* gene involved in DIM, but no PGL, biosynthesis (Simeone et al., 2010), displayed a lower mean ΔGP than the WT BCG strain (Figure 1b). In contrast, the BCGΔ*pks15/1* mutant, harbouring a mutation which blocks PGL synthesis without modifying DIM production (Constant et al., 2002), behaves similar to the WT BCG strain (Figure 1b). Together, our data indicate that DIM, and not PGL, contribute to the decrease in membrane polarity which extends around the site of bacterial contact over at least 1-1.5 μm. They also suggest that after transfer to the host membranes, DIM diffuse laterally in the membrane.

**Figure 1.**
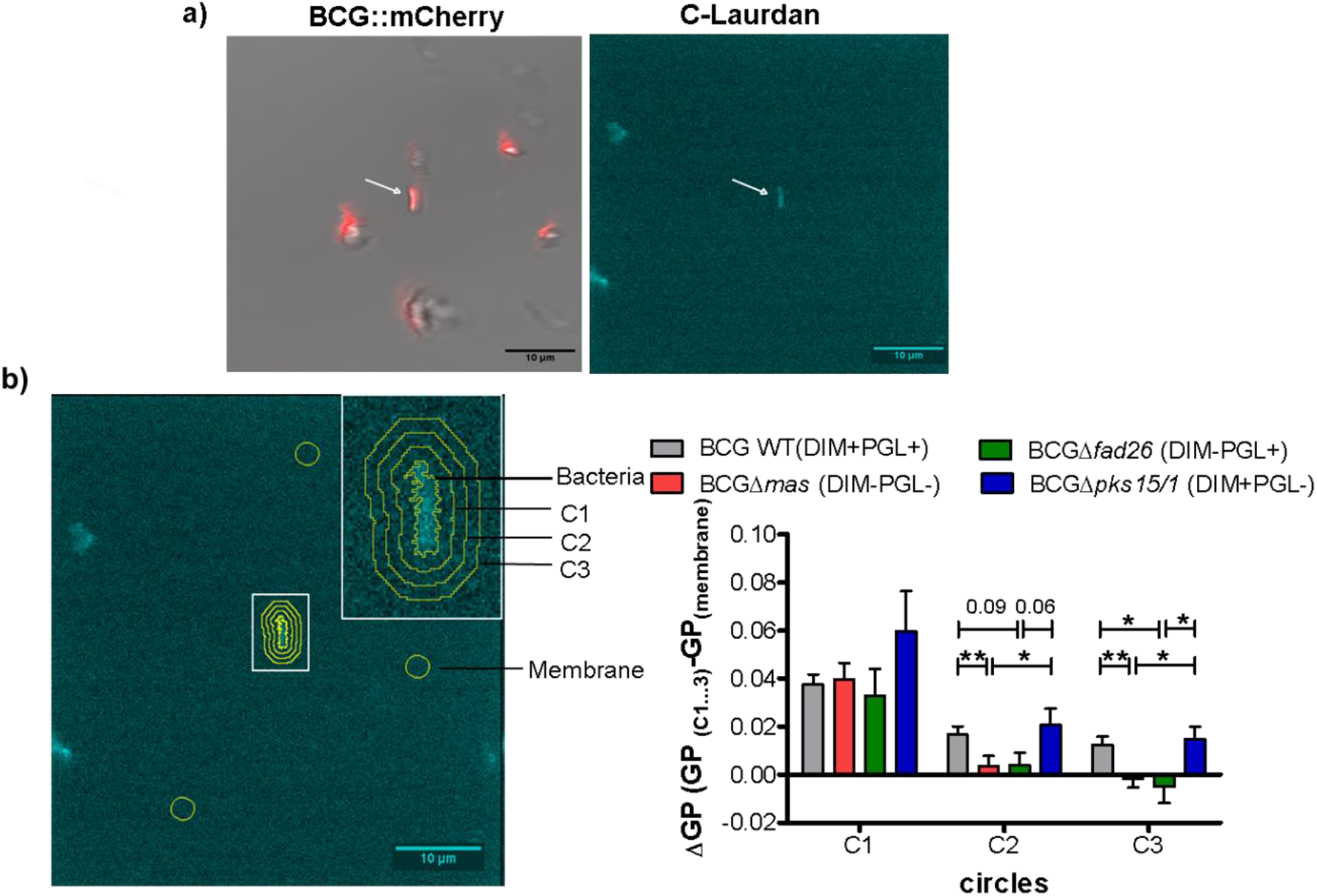
Analysis of DIM-induced changes in membrane polarity of supported bilayers. A POPC bilayer labelled with C-Laurdan was formed on a glass coverslip and incubated with 2 × 10^6^ *M. bovis* BCG::mCherry or BCGΔ*mas*::mCherry for 20 min. Two-photon microscopy images were taken on an inverted confocal microscope equipped with a femtosecond pulsed laser (Chameleon Vision II) and with a 561 nm DPSS laser for mCherry fluorescence and transmission imaging. A build-in spectrometer allowed the acquisition of a fluorescence spectrum in each pixel. C-Laurdan was excited at 720 nm; its fluorescence spectrum was collected in 18 channels between 418 nm and 593 nm (channel width 9.7 nm), resulting in a λ-stack of 18 images. The acquisition time per image was 31s **(a)** Immobilized bacteria were first selected by mCherry and transmission microscopy (left panel), then the C-Laurdan spectrum was acquired around an immobile bacterium by two-photon microscopy (right panel). **(b)** Contour and concentric bands around an immobile bacterium. ΔGP was calculated in these bands using the average GP of three random zones as the reference GP value of the membrane on each picture (left panel). Histograms represent the mean + SEM of 8 to 45 bacteria from 4 independent experiments (see SI Figure S1) (right panel). The statistical significance of difference in the ΔGP between strains was determined using Kruskal-Wallis’ test followed by Mann-Whitney’s test; * p< 0.05, **p<0.01.

### DIM induce the activation of complement receptor 3 for an optimal invasion of macrophages

We then asked whether this DIM-induced membrane perturbation could affect the phagocytic activity of CR3, the main receptor for *Mtb* entry in human macrophages. To address this question, we used zymosan, a yeast-derived polysaccharide particle, as a phagocytic prey for CR3 (Le Cabec et al., 2000). Macrophages were pre-incubated with exogenously added DIM or with the corresponding solvent. We then evaluated the effect of the CR3-specific blocking antibody 2LPM19c on the ability of macrophages to internalize zymosan (Figure 2a). In the absence of DIM, zymosan uptake was not significantly affected by adding 10μg/mL 2LPM19c, indicating that the CR3-dependent phagocytosis of zymosan is poorly efficient in resting macrophages (Figure 2a). Treatment of macrophages with DIM increased zymosan uptake by around 50% above the level without DIM (Figure 2a). We previously related this augmentation to the conical shape of DIM (Augenstreich et al., 2019). Here, we found that this DIM-induced increased was almost completely blocked by adding 2LPM19c (Figure 2a), supporting the proposal that DIM promote the engagement of CR3 in the phagocytosis of zymosan.

**Figure 2.**
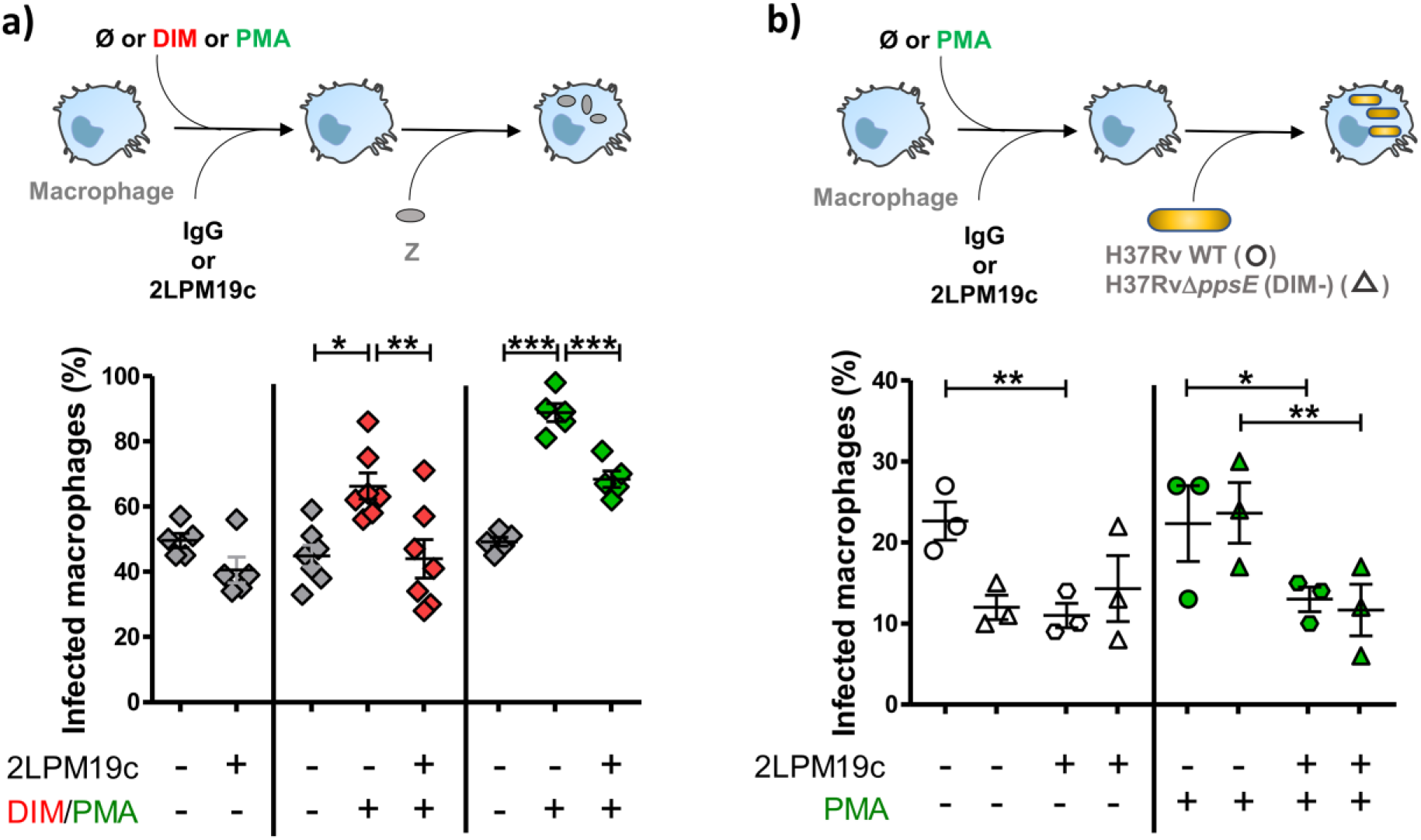
DIM and PMA trigger the entry of the DIM-deficient *H37Rv* mutant and of zymosan into macrophages through a CR3-dependent process. **(a)** Macrophages were either left untreated (grey square) or treated at 37°C for 1 h with lipid solvent (Ø, grey square) or 70 μM DIM (red square), or for 15 min with PMA solvent (Ø, grey square) or 50 nM PMA (green square). Cells were then incubated for a further 30 min with non-relevant IgG1 or 10 μg/mL 2LPM19c and put in contact for 1h with zymosan at MOI 30:1. **(b)** Macrophages were successively incubated with solvent (Ø, empty symbols) or 50 nM PMA (green symbols) and with IgG1 or 10 μg/mL 2LPM19c and then exposed to GFP-expressing H37Rv (circle) or H37RvΔ*ppsE* (triangle) at MOI 10:1 for 1 h. **(a, b)** At the end of infection, cells were rinsed, fixed and processed for the quantification of infected macrophages using a Leica 43 DM-RB epifluorescence microscope. For each set of conditions, the experiments were performed in duplicate, and at least 100 cells were counted per slide. The percentage of cells having ingested at least one bacterium, or one particle, was determined. The values are mean + SEM of 3-7 separate experiments. The significance of difference between control and treatment was evaluated using one-way ANOVA (a) or repeated measure ANOVA (b) followed by Bonferroni’s multiple comparison test; *p≤0.05, **p≤0.01, ***p≤0.001.

We then examined whether DIM affect surface expression of CR3. Using flow cytometry, we found no difference in CD11b expression between untreated macrophages and macrophages treated with DIM (Figure S2a). The functionality of CR3 also depends on the activation state of the receptor (Wright and Silverstein, 1982). Treating the macrophages with 50 nM phorbol myristate acetate (PMA), a known agonist activating CR3 (Diamond and Springer, 1993), increased phagocytosis of zymosan compared to untreated control cells. This effect is reduced by roughly 50% in the presence of 2LPM19c. Thus, the effect of purified DIM on phagocytosis resembles that of PMA (Figure 2a), supporting the hypothesis that DIM activate CR3 and facilitate zymosan phagocytosis.

Finally, we looked at what happens in the context of *Mtb*. As previously observed, 2LPM19c decreased by around 50 % the internalization of H37Rv WT, confirming that *Mtb* uses mainly CR3 to invade human macrophages (Astarie-Dequeker et al., 2009). Interestingly, adding PMA did not improve the uptake of H37Rv WT, suggesting that CR3 is already in an activated state (Figure 2b). In contrast, PMA restored both the infectivity of the DIM-deficient mutant H37Rv *ΔppsE*, deleted of *ppsE* encoding a protein essential for DIM biosynthesis and its inhibition by 2LPM19c to a level almost identical to that of the H37Rv WT strain (Figure 2b). Moreover, the CBRM1/5 antibody directed against the activation epitope of CR3 tended to block the uptake of both H37Rv both in untreated and PMA-treated cells and of H37RvΔ*ppsE* in PMA-treated cells only (Figure S2b). Together, these data led us to conclude that DIM activate CR3, thereby ensuring an optimal invasion of macrophages.

### DIM act in concert with mycobacterial effectors to promote lysis of biological membranes

We next enquired how DIM contribute to the membrane-disrupting effect of *Mtb*. Using our assay based on calcein leakage from large unilamellar vesicles (LUV) and mass spectrometry, we corroborated that the lytic activity of the batch of rEsxA from BEI, which we previously attributed to EsxA (Augenstreich et al., 2017), was in fact due to residual detergent ASB14 (Figure S3a-d) (Conrad et al., 2017). These observations prompted us to test a native version of EsxA (nEsxA) purified from a culture of *Mtb* (de Jonge et al., 2007). As expected, we observed pH-dependent lytic activity of nEsxA on LUV made of POPC (Figure 3a) (de Jonge et al., 2007). In addition, the activity at pH 5 was significantly decreased after digestion by proteinase K, ruling out the presence of detergent (Figure 3a). Based on these positive results, we investigated whether DIM could modulate the activity of nEsxA. On liposomes of THP-1 lipids without DIM, the lytic activity of nEsxA at pH 5 was less than 10% (Figure 3b), well below that observed on POPC liposomes (Figure 3a). This difference can be explained by the higher membrane rigidity of THP-1 liposomes compared to POPC liposomes (Augenstreich et al., 2017;Ray et al., 2019). Nevertheless, incorporating DIM in THP-1 liposomes tended to potentiate by 35 % the activity of nEsxA (Figure 3b). We also tested a recombinant version of EsxA (rEsxA) (Figure 3b) produced by us in *E. coli* and purified using the protocol provided by BEI Resources (Conrad et al., 2017) without the ASB-14-based endotoxin removal. We noticed that rESXA permeabilized liposomes more efficiently than nEsxA. We have no rational explanation for this difference but we observed that DIM also increased the membranolytic activity of rEsxA (Figure 3b). This effect is specific to DIM, as the incorporation of the apolar triglyceride tripalmitin in THP-1 liposomes had no impact on the calcein leakage induced by rESXA (60+30 % and 53+15 %, n=2, in the absence and the presence of 10 % tripalmitin, respectively). We also examined the effect of DIM on other lytic agents characterized by distinct structures: the detergent ASB-14 and the two membrane-disrupting proteins melittin and perforin-1. Incorporation of DIM into THP-1-derived liposomes increased the lytic activity of all three agents (Figure S3e and Figure 3b). These observations indicated that DIM enhance the activity of different membranolytic compounds with different structures and modes of action.

**Figure 3.**
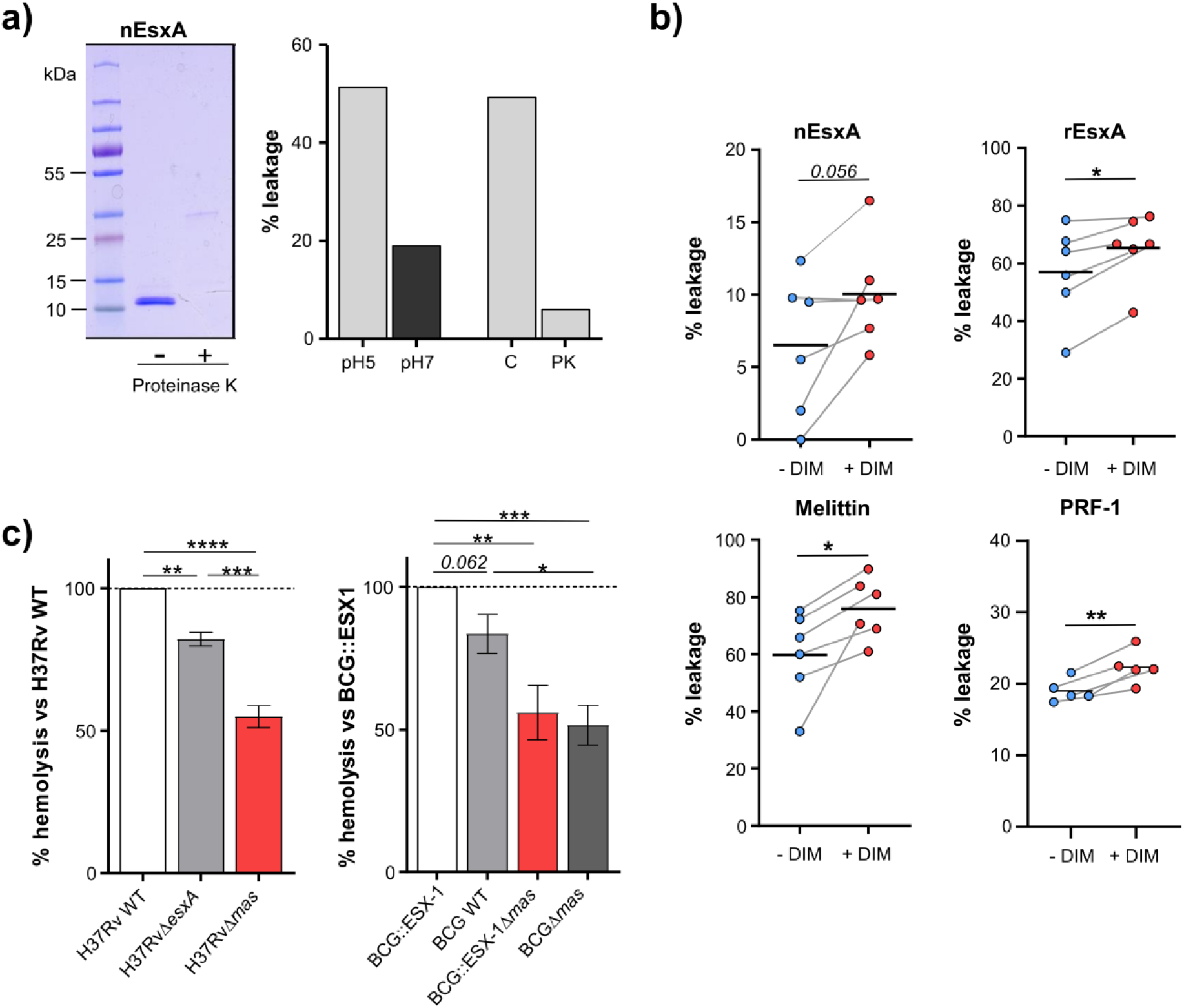
DIM potentiate EsxA membranolytic activity *in vitro*, and is required for EsxA hemolytic activity. **(a)** left panel: the active form of the native EsxA (nEsxA) untreated (-) or digested with proteinase K (+) was analyzed by SDS-PAGE followed by Coomassie Blue staining; right panel: POPC liposomes were put in contact with 10 μM nEsxA at pH 5 or pH 7, or with 10 μM intact or proteinase K-digested nEsxA at pH5. **(b)** Calcein leakage in THP-1 liposomes supplemented or not with 10% of DIM (w/w) and incubated with 10 μM nEsxA (pH 5), rEsxA (pH 5), 50 nM mellitin (pH 7) or 15 nM perforin-1 (PRF-1) (pH 7). **(c)** Strains of H37Rv or BCG producing a combination of EsxA and DIM were put in contact for 48 h at 37°C with 1 × 10^7^ erythrocytes at a MOI 50:1. The supernatant was collected and the OD_415_ was measured and normalized to OD_415_ of the corresponding DIM- and EsxA-proficient strain. The values are mean + SEM of 4-5 separate experiments. The significance of difference in the percentage of leakage between untreated and membranolytic agent-treated erythrocytes was evaluated using the paired Student’s t test. The significance of difference between strains was determined using a one-way ANOVA test followed by Bonferroni’s multiple comparison test; *p≤0.05, **p≤0.01, ***p≤0.001; ****p≤0.0001.

We also tested whether DIM work with EsxA in a mycobacterial context. To this end, we studied the hemolytic activity of several knock-out or knock-in mutants of *Mtb* and *M. bovis* BCG expressing combinations of DIM and EsxA. These strains have already been characterized for their capacity to produce EsxA and DIM (Augenstreich et al., 2017). The H37Rv WT strain incubated for 48 h with human erythrocytes exhibited hemolytic activity (Figure 3c). The H37RvΔ*esxA* and H37RvΔ*mas* mutants were significantly less hemolytic than the H37Rv WT strain (Figure 3c), showing that both DIM and EsxA contribute to hemolysis. These results were confirmed in the BCG background, which has a partial deletion of ESX-1 and a functional inhibition of EsxA. Indeed, when BCG was genetically complemented with the ESX-1 region from *Mtb*, EsxA secretion was re-established (Augenstreich et al., 2017). The resulting BCG:: ESX-1 strain demonstrated higher hemolytic activity than the BCG WT strain and the DIM-deficient BCG::ESX-1Δ*mas* mutant (Figure 3c). Most notably, in both genetic backgrounds, the DIM-deficient strains exhibited significantly lower hemolytic activity than EsxA- or ESX-1-deficient strains, showing that DIM have an impact extending beyond EsxA/ESX-1.

## Discussion

The notion of DIM as bacterial effectors subverting the host’s immune responses has recently emerged (Rousseau et al., 2004;Astarie-Dequeker et al., 2009;Cambier et al., 2014;Passemar et al., 2014;Augenstreich et al., 2017;Barczak et al., 2017;Conrad et al., 2017;Quigley et al., 2017;Lerner et al., 2018), but the molecular mechanisms by which DIM exert their biological activity are poorly known. We have previously demonstrated that DIM are transferred from the envelope of *Mtb* to the membranes of host macrophages during infection and locally disturb their lipid organization (Astarie-Dequeker et al., 2009;Augenstreich et al., 2019). The next step was to identify the functional role of this transfer. We now establish that DIM induce a membrane perturbation, which spreads beyond the point of contact with the bacilli and improves the activity of proteins either embedded in host membranes, like CR3, or acting on these membranes, like EsxA and other membranolytic effectors.

Using C-laurdan combined with two-photon microscopy, we showed that the membrane polarity decreased at the site of interaction of BCG with a lipid membrane. This is consistent with our previous observation of a global decrease in membrane polarity in a population of macrophages infected by BCG (Astarie-Dequeker et al., 2009). As C-Laurdan allows identifying lipid domains with different packing degrees and lipid order (Owen et al., 2010), the observed decrease in polarity strongly supports the proposal that DIM locally disturb the lipid organization of host membranes and alter their physical properties. This can also be interpreted in the light of our recent findings showing that DIM adopt a conical molecular shape in a simple phospholipid bilayer and can disorganize such a bilayer by promoting the formation of non-bilayer (inverted hexagonal) membrane phases (Augenstreich et al., 2019). Remarkably, we observed that the local decrease in membrane polarity extends over a distance of 1 to 1.5 μm around the bacterium. These observations provide experimental support to our molecular dynamic simulations predicting diffusion of DIM inside the membrane (see (Augenstreich et al., 2019), supplementary movie). Indeed, after transfer from the bacterium to the target membranes, DIM diffuse away from the contact site, causing a gradual fading of the polarity.

Changes in membrane lipid composition and organization are generally thought to influence many cellular functions. Our findings go beyond the functional observations that *Mtb* uses envelope lipids to remodel the host membrane and divert macrophages functions (Laneelle and Tocanne, 1980;Sut et al., 1990;Welin et al., 2008;Augenstreich et al., 2019;Mishra et al., 2019). They provide evidence that insertion of lipids into host cell membranes helps the bacteria exploit membrane partners that direct the cell’s responses. For instance, DIM increase CR3 activity for promoting phagocytosis. Like most integrin, CR3 is normally exposed at the cell surface in a low-activity state but can undergo changes that lead to increased activity, *e.g*. conformational changes leading to a state of higher affinity for their specific ligands (Liddington and Ginsberg, 2002). We found that DIM mimic at least in part the action of PMA, a known activator of CR3. For instance, DIM treatment increased in the uptake of zymosan, which was almost completely abolished when CR3 was blocked by an anti-CR3 blocking Antibody. Moreover, PMA restored the ability of a DIM-deficient mutant to infect macrophages equally efficiently as the WT strain, probably through the engagement of the active epitope of CR3 as assessed using CBRM1/5. All together, these data tend to indicate that DIM can also activate the CR3 receptor. This activation would enable recognition by CR3 of PAMPs at the surface of *Mtb* leading to its uptake. Conical lipids like DIM may induce membrane curvature (Cooke and Deserno, 2006). The resulting membrane elastic stress may in turn modulate the function of integral membrane proteins (Phillips et al., 2009). It is therefore reasonable to hypothesize that DIM impose a curvature on the host’s membrane that governs CR3’s activation. This is consistent with the transmembrane nature of CR3 and its lateral mobility within cell membranes (Ross et al., 1992). DIM could also act on other membrane receptors cooperating with CR3 to carry out entry of mycobacteria, like CD14 which has been shown to induce an inside-out signaling pathway involving TLR2, leading to a CR3-dependent phagocytosis of *M. bovis* BCG (Sendide et al., 2005).

Our findings reinforce the proposal that DIM can work with *Mtb*EsxA (Augenstreich et al., 2017;Barczak et al., 2017). Using liposomes formed of THP-1 lipids, we demonstrate that DIM enhance the activity of *Mtb* EsxA. Combined with our previous data in macrophages, these results provide a reasonable explanation for DIM’s involvement in the deterioration of the phagosomal membrane by *Mtb* (Augenstreich et al., 2017;Quigley et al., 2017). Interestingly, this potentiating effect of DIM is not specific to EsxA but also observed for the two pore-forming proteins melittin and perforin-1, and even for the detergent ASB-14. This strengthens the evidence that DIM act by perturbing their target membrane to promote the membrane-disruptive activities of these molecules. Furthermore, using gene-deleted *Mtb* mutants, we observed that the loss of DIM results in a lower hemolytic activity of *Mtb* than the loss of EsxA. As DIM alone have no lytic activity, this implies that additional bacterial factors are involved in this process.

In conclusion, our findings reveal molecular mechanisms by which *Mtb* exploit DIM to rewire the host cells. Given their overall hydrophobic properties, DIM are able to reshape locally the host cell membrane in a way that targets cell signaling pathways and bacterial effectors, thus supporting their multifaceted role in the virulence of *Mtb*.

## Supporting information

Supplemental Informations

## Acknowledgments

This work was supported by the European Union’s Horizon 2020 Research and Innovation Program (TBVAC2020, 643381), the Agence Nationale de la Recherche (ANR-10-LABX-62-IBEID, ANR-14-JAMR-001-02 and ANR-16-CE15-0003), the Fondation pour la Recherche Médicale FRM (DEQ20130326471 and DEQ20160334879) and the Centre National de la Recherche Scientifique (CNRS). J.A. was a recipient of a PhD scholarship from the French government. C.A.D. and C.G thank the TRI-Genotoul Imaging facility (Toulouse, France) and S. Dauvillier for technical help and fruitful discussions.

