## Supplemental Informations for "Phthiocerol dimycocerosates from *Mycobacterium tuberculosis* increase the membrane activity of bacterial effectors and host receptors"

This file includes:

- Figures and legends S1 to S3

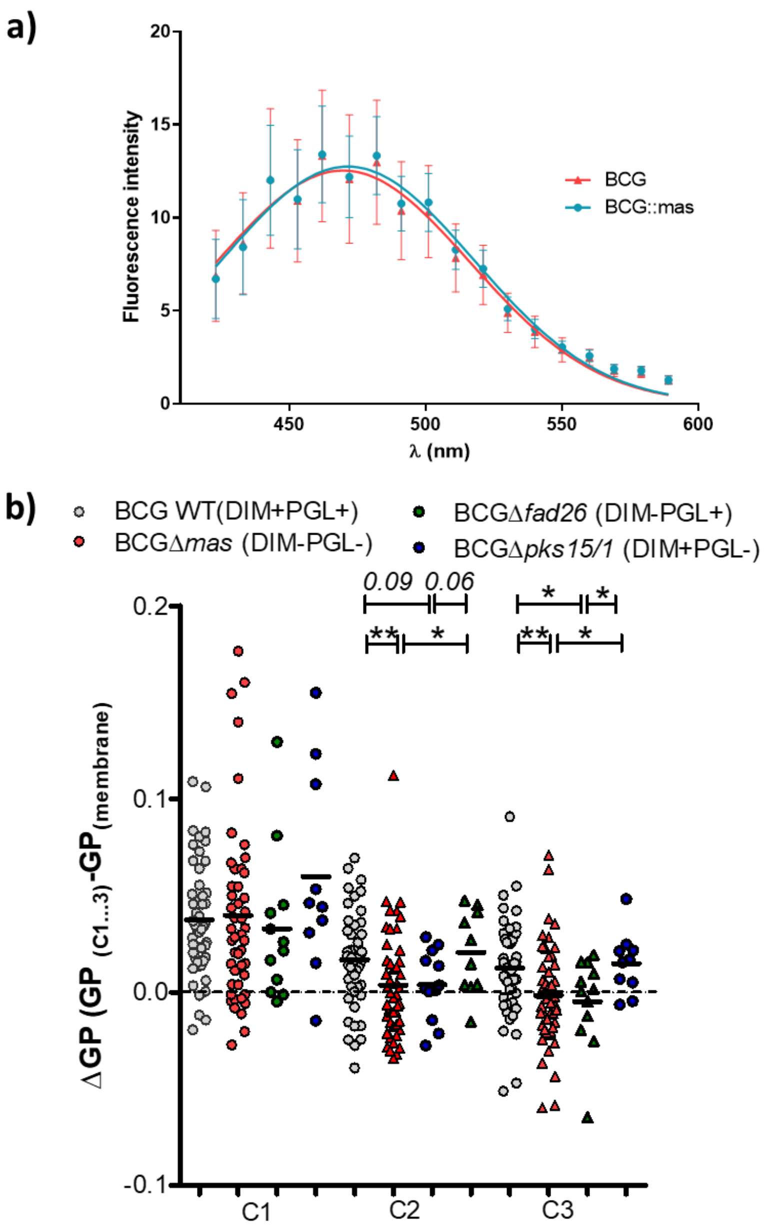


**Figure S1. Analysis of DIM-induced changes in membrane polarity of supported bilayers put in contact with BCG strains. (a)** *M. bovis* BCG::mCherry or BCG*Δmas*::mCherry were labelled with C-Laurdan, excited at 720 nm and the fluorescence spectrum of c-Laurdan was collected for each strains in 18 channels ranging from 418 nm to 593 nm (channel width 9.7 nm), resulting in a stack of 18 images (λ-stack). **(b)** A POPC bilayer labelled with C-Laurdan was formed on a glass coverslip and incubated with $2x10^6$ *M. bovis* BCG::mCherry or BCG*Δmas*::mCherry for 20 min. Adherent bacteria were selected and the fluorescence spectrum was acquired by two-photon microscopy. The ΔGP was calculated as indicated in the legend of Figure 1. Each symbol within vertical scatter plots represents a ΔGP for one bacterium; The mean value is represented by the dark grey line. The statistical significance of difference in the ΔGP between strains was determined using a Kruskal-Wallis' test followed by a Mann-Whitney's test, * $p< 0.05$, ** $p<0.01$.

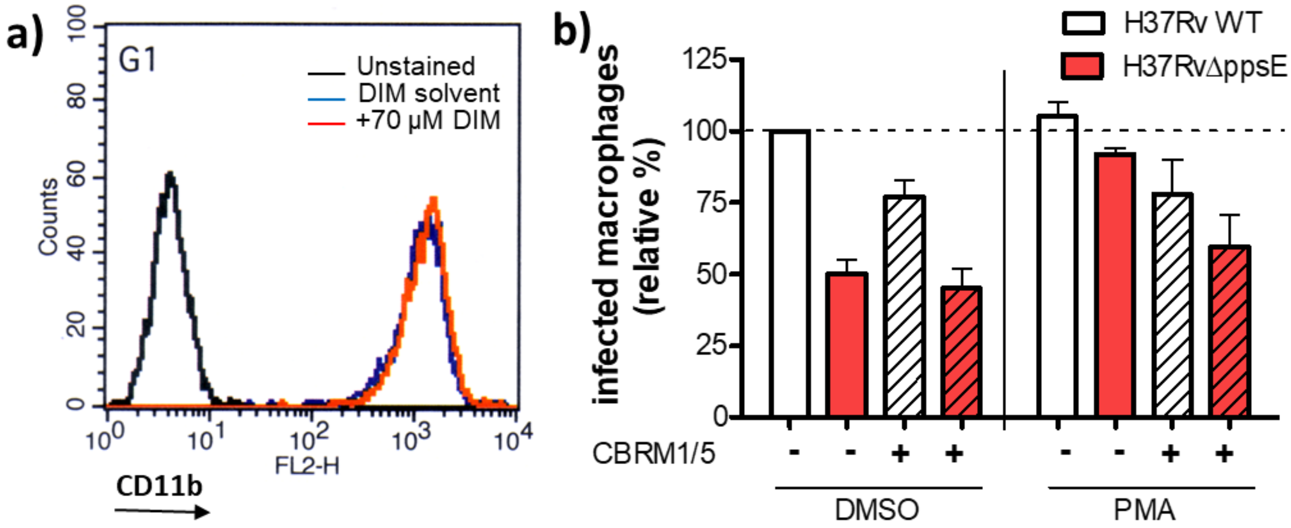


**Figure S2. (a) DIM treatment does not affect CR3 expression at the surface of macrophages.** Macrophages were treated with lipid solvent (blue line) or 70 µM DIM (red line), incubated for 1 h at 37°C and processed for evaluation of membrane-bound CR3 expression by flow cytometry. Briefly, cells were stained with mAbCD11b-PE (BD Biosciences, San Jose, CA) (blue and red lines) or the corresponding isotype control antibody (black line). Flow cytometry was performed using LSR-II flow cytometer (BD Biosciences) and the associated BD FACSDiva softwareLSR II analyzer (BD Bioscience and data were analyzed using FlowJo software (FlowJo). **(b) Blocking the activation epitope of CR3 decreases the uptake of both H37Rv and H37RvΔ*ppsE* in PMA-treated cells**. Macrophages were successively incubated with PMA solvent (DMSO) or 50 nM PMA for 15 min and with the non-relevant IgG1 or 10 µg/mL anti-CR3 mAb CBRM1/5 directed against the activation epitope of CR3 for 30 min. Cells were then exposed for 1h at 37°C to GFP-expressing H37Rv (White bar) or H37RvΔ*ppsE* (red bar) at MOI 10:1. The histogram represents the percentage of macrophages infected with H37Rv and H37RvΔ*ppsE* in treated and untreated macrophages, expressed with respect to H37Rv WT in untreated cells (100%). The values are means ± SEM of 2 separate experiments.

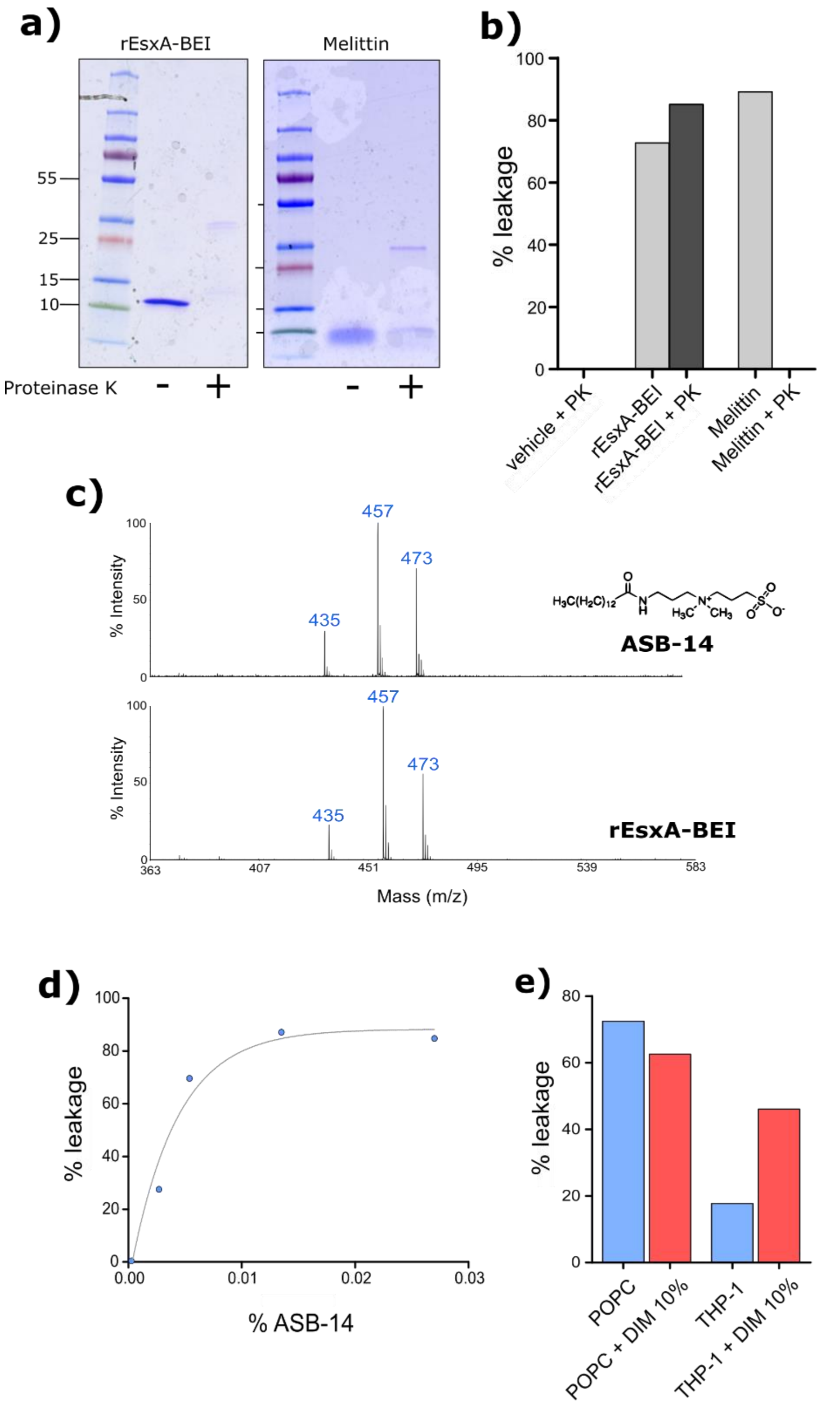


**Figure S3. The detergent ASB-14 is detected in rEsxA-BEI samples and exerts a membranolytic activity that can be modulated by DIM. (a)** 10 µg of rEsxA and Mellitin were digested by proteinase K for 1h at 37°C. The digestion was verified by SDS-PAGE and Coomassie Blue staining. **(b)** The activity of intact or digested rEsxA (10 µM) and Melittin (50 nM) was tested on POPC liposomes at pH7. **(c)** 2.5 µg of ASB-14 and 1 µg of EsxA were analyzed by mass spectrometry MALDI-TOF. The mass spectrum show three peaks representing ASB-14 ions at 435 Da ($M+H^+$) and pseudomolecular ion $(M+Na)^+$ and $(M+K)^+$ at 457 Da and 473 Da respectively**. (d)** The membranolytic activity of ASB-14 was tested at different doses on POPC liposomes using calcein leakage assay. **(e)** The membranolytic activity of 0.05% ASB-14 was tested on POPC liposomes and THP-1 liposomes supplemented or not with 10% DIM (w/w).
